# Longitudinal single-cell profiling links RUNX1T1 to lineage transition and maintenance in treatment-induced neuroendocrine prostate cancer

**DOI:** 10.1101/2025.05.14.653660

**Authors:** Yuchao Ni, Mingchen Shi, Dong Lin, Yen-Yi Lin, Hui Xue, Xin Dong, Liangliang Liu, Funda Sar, Rebecca Wu, Tunc Morova, Anne Haegert, Robert Bell, Xinyao Pang, Adam Classen, Wei Dong, Yu Wang, Junru Chen, Stéphane Le Bihan, Vickie Wang, Ning Xu, Nathan Lack, Martin Gleave, Christopher Ong, Gang Wang, Hao Zeng, Colin Collins, Yuzhuo Wang

## Abstract

Treatment-induced neuroendocrine prostate cancer (t-NEPC) is a lethal form of advanced prostate cancer that can emerge from adenocarcinoma under androgen receptor pathway inhibition, but the tumor-cell states and regulators underlying this transition remain incompletely defined. Here, we analyzed longitudinal single-cell RNA sequencing data from the LTL331/331R patient-derived xenograft model across seven disease stages, comprising 32,269 tumor-cell transcriptomes from treatment-naive adenocarcinoma, post-castration regression, and relapsed NEPC. We identified an AR-^low^/NE-^low^ intermediate state that emerged after castration and before overt relapse, as well as two transcriptionally distinct terminal NEPC states marked by ASCL1^high^/FOXA2^low^ and ASCL1^low^/FOXA2^high^ programs. RUNX1T1 was induced in the intermediate state and remained elevated across both terminal states. RUNX1T1 enhanced NE-associated features and cell viability during enzalutamide treatment in adenocarcinoma cells, whereas its knockdown in NCI-H660 cells reduced NE-associated transcriptional programs, proliferation, and survival and shifted the transcriptome toward an intermediate-like state. Chromatin interactome analysis and co-immunoprecipitation linked RUNX1T1 to G9A/LASP1-containing chromatin repressor complexes. These findings define a temporal framework for treatment-induced NEPC progression and identify RUNX1T1 as an early-induced and sustained regulator of NE lineage transition and maintenance of established NEPC.

## Introduction

Treatment-induced neuroendocrine prostate cancer (t-NEPC) is a highly aggressive form of advanced prostate cancer that can arise through lineage plasticity involving transdifferentiation of prostate adenocarcinoma under the selective pressure of androgen deprivation therapy (ADT) and potent androgen receptor pathway inhibitors (ARPIs) [1–5]. t-NEPC is characterized by reduced androgen receptor (AR) signaling, acquisition of neuroendocrine (NE) features, and resistance to AR-directed therapies. Patients with t-NEPC have poor clinical outcomes, underscoring the need to understand how this lineage transition develops and how the resulting NEPC state is maintained [6–10].

Recent studies have advanced our understanding of established NEPC, including its lineage programs, aggressive behavior, epigenetic regulation, and therapeutic vulnerabilities [1, 11–17]. Longitudinal studies in genetically engineered mouse models have also provided important insight into NEPC evolution [18, 19]. However, the temporal evolution of tumor-cell states during treatment-induced adenocarcinoma-to-NEPC transdifferentiation in human prostate cancer remains less clear, particularly the relationship between early post-treatment reprogramming and the cellular states present at relapse. Mapping this process from early post-treatment reprogramming through relapse may define cell-state evolution and identify regulators induced early and retained after NEPC is established, a temporal pattern consistent with roles in both the acquisition and maintenance of NE-associated features.

To address this gap, we used the longitudinal LTL331/331R patient-derived xenograft (PDX) model of treatment-induced adenocarcinoma-to-NEPC transdifferentiation, which recapitulates the donor patient’s clinical course and provides a clinically relevant platform for studying t-NEPC evolution [20–22]. Previous integrated genomic and bulk transcriptomic analyses of this model showed that t-NEPC arises through adaptive transcriptional reprogramming [22]. Using this system, we and our collaborators have identified multiple regulators implicated in t-NEPC development and progression, including HP1α, ONECUT2, BRN2, PEG10, SRRM4, PROX1, PRC2, and the CXCR4-LASP1-G9A-SNAIL repressive axis [13–15, 22–27]. These studies establish LTL331/331R as a well-characterized model for dissecting the regulatory architecture of treatment-induced NE lineage plasticity.

Here, we analyzed longitudinal single-cell RNA sequencing data from the LTL331/331R model to examine tumor-cell states across treatment-induced adenocarcinoma-to-NEPC progression. We identified an intermediate state that emerged after castration and before fully relapsed NEPC, together with two transcriptionally distinct terminal NEPC states marked by ASCL1^high^ and FOXA2^high^ programs. RUNX1T1 was induced in the intermediate state and remained elevated across both terminal NEPC states. Functional studies showed that RUNX1T1 enhanced ARPI-induced NE-associated features and increased cell viability during enzalutamide treatment in adenocarcinoma cells and supported NEPC identity, proliferation, and survival in NCI-H660 cells. Chromatin interactome analysis and co-immunoprecipitation further linked RUNX1T1 to G9A/LASP1-containing chromatin repressor complexes. Together, these findings identify RUNX1T1 as an early-induced and sustained regulator linking treatment-induced lineage transition to maintenance of established NEPC.

## Results

### 1. Longitudinal single-cell profiling of the LTL331/331R PDX model reveals three tumor-cell states during treatment-induced adenocarcinoma-to-NEPC transdifferentiation

To characterize tumor-cell states during treatment-induced NEPC transdifferentiation, we analyzed longitudinal scRNA-seq data from the LTL331/331R PDX model across seven disease stages spanning pre-castration adenocarcinoma, post-castration regression, and relapsed NEPC [27]. After quality control, 32,269 tumor-cell transcriptomes were retained for analysis (Supplementary Table S1).

Unsupervised clustering identified 17 transcriptionally distinct tumor-cell clusters organized into three major states based on AR activity and lineage-marker expression (Fig. 1A-E; Supplementary Table S2). The adenocarcinoma compartment included pre-castration AR^+^/PSA^+^/NE^-^clusters 10 and 15, which showed the highest AR activity, and post-castration AR^+^/PSA^low/-^/NE^-^clusters with progressively reduced AR activity (Fig. 1C-E). Two terminal NEPC clusters, clusters 11 and 13, displayed an AR^-^/PSA^-^/NE^+^ phenotype, with diminished AR activity and increased NE-marker expression (Fig. 1B-E). Cluster 16 exhibited an AR^low^/PSA^-^/NE^low^ phenotype, occupied an intermediate position between the adenocarcinoma and terminal NEPC populations, and showed the strongest EMT signature across the tumor-cell clusters (Fig. 1A,C,D). Notably, terminal NEPC cells retained expression of the luminal keratins KRT8 and KRT18, consistent with their emergence through lineage reprogramming of luminal adenocarcinoma cells.

**Figure 1.**
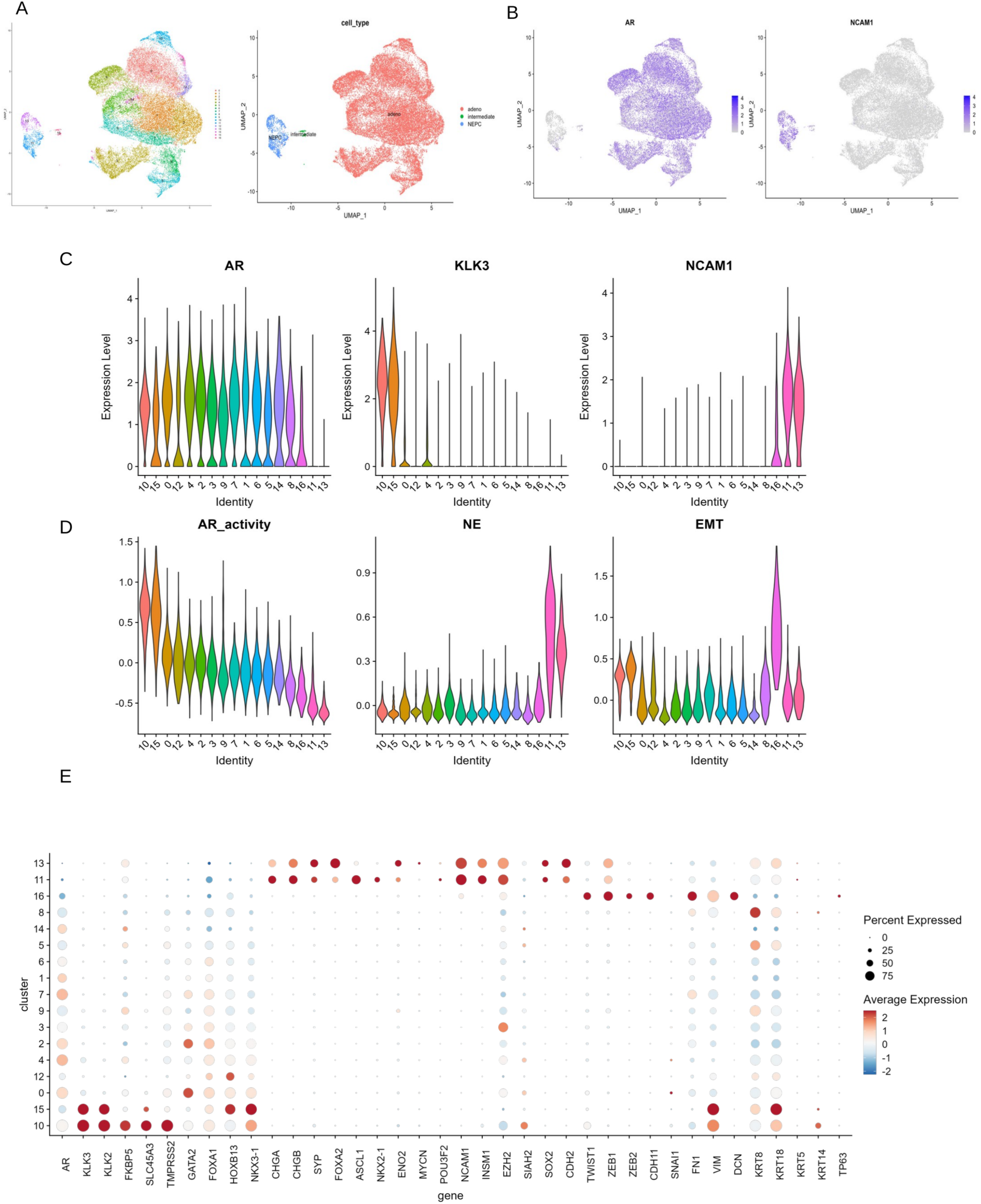
Longitudinal single-cell profiling resolves tumor-cell states during treatment-induced adenocarcinoma-to-NEPC progression. **(A)** UMAPs of 32,269 human tumor cells from seven LTL331/331R time-course samples, colored by 17 unsupervised clusters (left) or grouped into adenocarcinoma, intermediate, and terminal neuroendocrine prostate cancer (NEPC) states (right). Cluster 16 represents the intermediate state, and clusters 11 and 13 represent terminal NEPC. In the state-level UMAP, adenocarcinoma, intermediate, and terminal NEPC cells are shown in salmon, green, and blue, respectively. **(B)** Feature UMAPs showing log-normalized AR and NCAM1 expression. Gray indicates no detected expression on the plotted scale, and the purple-to-blue scale indicates increasing expression. **(C)** Violin plots of AR, KLK3, and NCAM1 expression across clusters. **(D)** Violin plots of androgen receptor (AR) activity, neuroendocrine (NE), and epithelial-mesenchymal transition (EMT) signature scores. In C and D, clusters are colored consistently and ordered from higher to lower AR activity. **(E)** Dot plot of selected lineage and cell-state markers. Dot color indicates average scaled expression, with red and blue representing higher and lower relative expression, respectively; dot size indicates the percentage of cells expressing each gene.

Together, these findings define three major states across the disease course: AR^+^/NE^-^adenocarcinoma with declining AR activity after castration, an AR^low^/NE^low^ intermediate state represented by cluster 16, and AR^-^/NE^+^ terminal NEPC represented by clusters 11 and 13.

### 2. Intermediate cluster 16 defines a putative transitional state during NE transdifferentiation

Having defined the major tumor-cell states, we next examined their temporal distribution and inferred relationships across the LTL331/331R time course. Terminal NEPC clusters were absent before castration and became prominent during late post-castration progression and relapse. Cluster 16 was also absent before castration but emerged during post-castration regression, before terminal NEPC became dominant (Fig. 2A,B; Supplementary Table S1). This temporal distribution placed cluster 16 early in the treatment-induced lineage transition.

**Figure 2.**
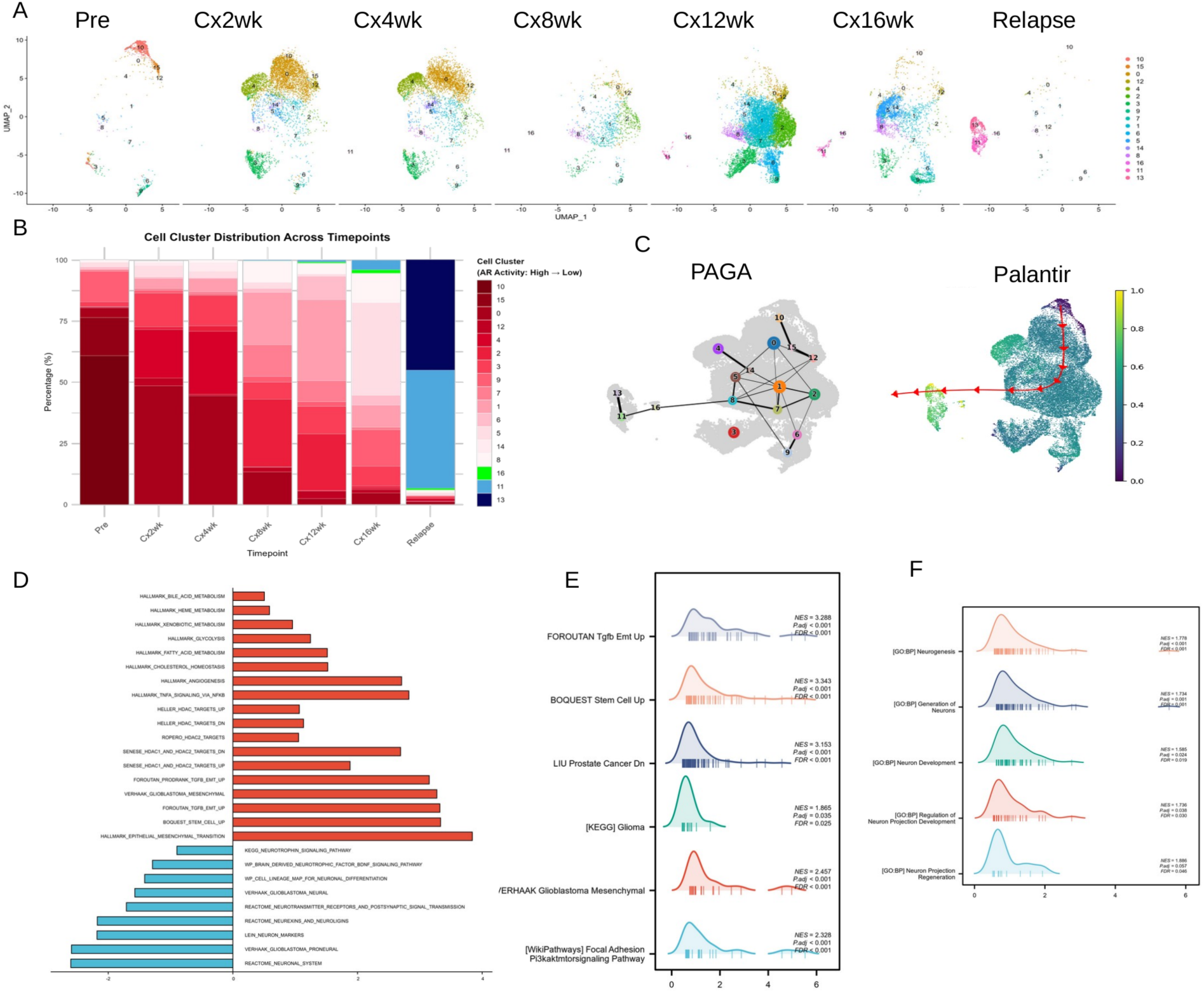
Longitudinal analysis identifies a putative intermediate state during treatment-induced NEPC progression. **(A)** UMAPs showing tumor-cell clusters at the pre-castration stage, 2, 4, 8, 12, and 16 weeks post-castration (post-Cx), and relapse. **(B)** Proportional distribution of clusters across the seven time points. Adenocarcinoma clusters are colored from red to pink according to decreasing AR activity, cluster 16 is shown in green, and terminal NEPC clusters 11 and 13 are shown in light and dark blue, respectively. **(C)** Partition-based graph abstraction (PAGA) connectivity and Palantir pseudotime analyses. In the PAGA graph, numbered nodes represent clusters, edges represent inferred connectivity, and edge thickness represents connection strength. In the Palantir map, color indicates relative pseudotime from 0 to 1. **(D)** Gene set enrichment analysis (GSEA) comparing cluster 16 with terminal NEPC clusters. Red bars and positive normalized enrichment scores (NES) indicate programs enriched in cluster 16, whereas blue bars and negative NES indicate programs enriched in terminal NEPC. Selected lineage-plasticity, metabolic, chromatin-regulatory, and neuronal programs are shown. **(E)** Representative enrichment profiles for EMT, TGF-β-associated EMT, mesenchymal, stem cell-associated, and related lineage-plasticity programs enriched in cluster 16. **(F)** Representative enrichment profiles for neurogenesis and neuronal-development programs enriched in terminal NEPC relative to cluster 16. In E and F, the colored curve represents the running enrichment score, vertical bars indicate positions of gene-set members in the ranked list, and NES, P values, and false-discovery rates (FDRs) are shown.

Trajectory analysis using PAGA and Palantir independently positioned cluster 16 between the adenocarcinoma and terminal NEPC populations (Fig. 2C). The agreement between its temporal emergence and inferred trajectory position supports classification of cluster 16 as a putative transitional state during NE lineage progression.

Pathway enrichment analysis further defined the transcriptional features of cluster 16 (Fig. 2D-F). Consistent with the EMT-enriched differentiation node previously reported in this model [27], cluster 16 showed strong enrichment for TGF-β-driven EMT, mesenchymal-like programs, and stem cell-associated signatures, consistent with features of lineage plasticity. Adenocarcinoma-associated programs were reduced relative to adenocarcinoma clusters, whereas neurodevelopmental and NE-associated programs remained less active than in terminal NEPC. Thus, cluster 16 had attenuated adenocarcinoma identity without fully acquiring the terminal NEPC program.

Cluster 16 was also enriched for glycolysis, fatty acid metabolism, cholesterol homeostasis, bile acid metabolism, and xenobiotic metabolism (Fig. 2D). HDAC-associated signatures were enriched as well, suggesting that metabolic and chromatin-regulatory programs accompany this intermediate state. Together, its temporal distribution, trajectory position, and transcriptional features support cluster 16 as a putative transitional state with features of lineage plasticity and enrichment of metabolic and chromatin-regulatory programs.

### 3. Distinct ASCL1^high^ and FOXA2^high^ states reveal dynamic heterogeneity within terminal NEPC

We next asked whether the two terminal NEPC clusters identified above, clusters 11 and 13, represented distinct transcriptional states. Given the established role of ASCL1 in NE differentiation across multiple NE cancers, including small cell lung cancer (SCLC) and NEPC, we first examined ASCL1 expression across the two NEPC clusters. Cluster 11 showed high ASCL1 expression, whereas cluster 13 exhibited lower ASCL1 expression together with a distinct transcriptional profile (Fig. 3A).

**Figure 3.**
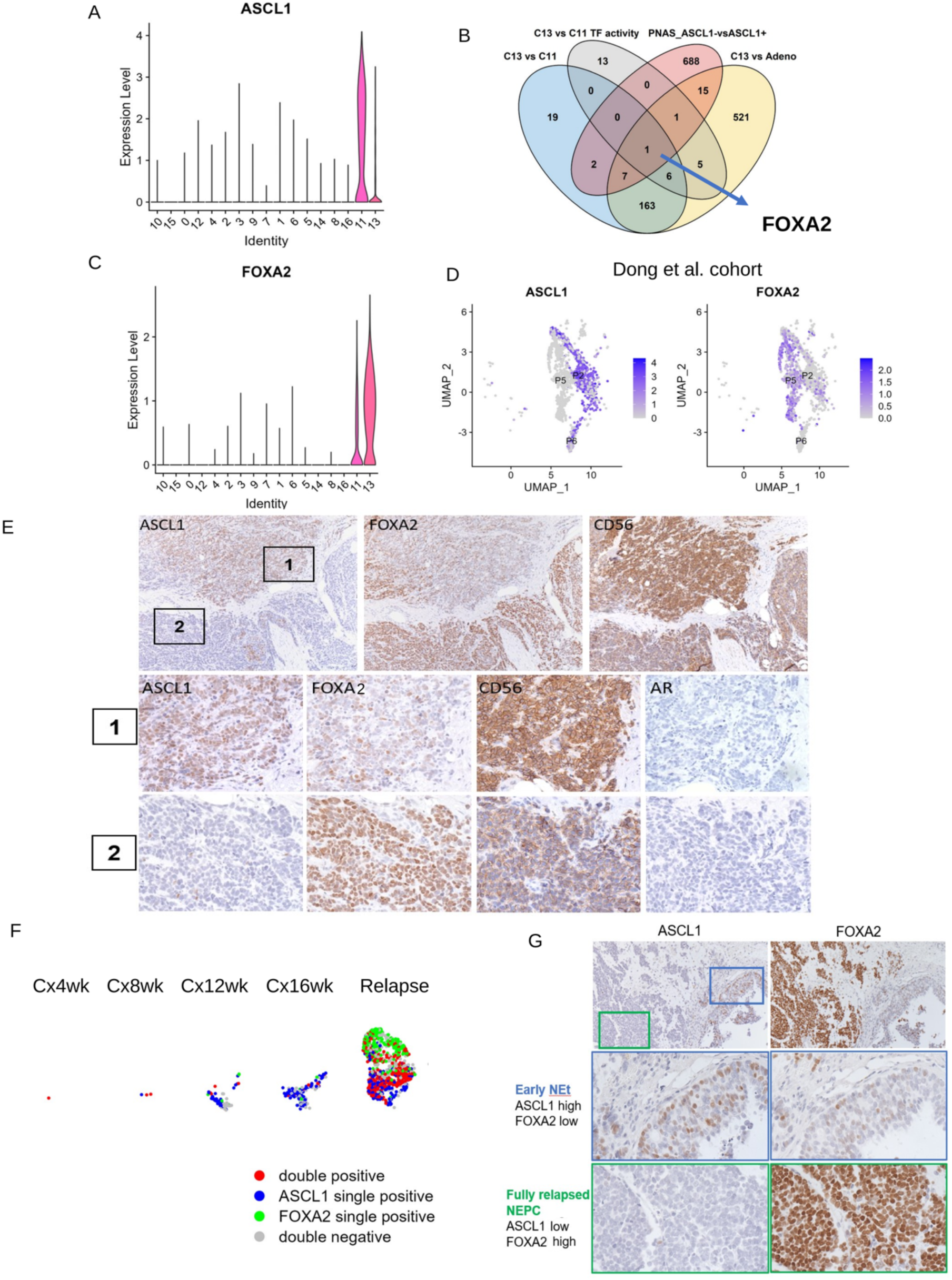
Distinct ASCL1-high and FOXA2-high states reveal dynamic heterogeneity within terminal NEPC. **(A)** Violin plot of ASCL1 expression across tumor-cell clusters, showing preferential expression in terminal NEPC cluster 11. **(B)** Four-set intersection used to identify transcription factors with higher expression in cluster 13 than in adenocarcinoma clusters, higher expression in cluster 13 than in cluster 11, higher inferred activity in cluster 13 than in cluster 11, and concordant upregulation in clinical ASCL1^low^ versus ASCL1^high^ NEPC cells. FOXA2 was the only transcription factor meeting all four criteria. **(C)** Violin plot of FOXA2 expression across tumor-cell clusters, showing higher expression in terminal NEPC cluster 13 than in cluster 11. **(D)** Feature UMAPs showing normalized ASCL1 and FOXA2 expression in a published clinical NEPC single-cell RNA-sequencing cohort [29]. Gray indicates zero plotted expression, and the purple-to-blue scale indicates increasing expression. **(E)** Representative immunohistochemistry (IHC) from a clinical NEPC tumor. Low-magnification serial sections identify regions 1 and 2; corresponding higher-magnification images show relative ASCL1^high^/FOXA2^low^ and ASCL1^low^/FOXA2^high^ patterns, respectively. CD56 and AR staining provide phenotypic context. Corresponding spatial patterns were identified in 9 of 10 clinical NEPC specimens. **(F)** Temporal distribution of ASCL1^+^/FOXA2^+^ (red), ASCL1^+^/FOXA2^-^(blue), ASCL1^-^/FOXA2^+^ (green), and double-negative (gray) cells at the indicated post-castration time points and relapse. **(G)** Representative ASCL1 and FOXA2 IHC during early NE transition and fully relapsed NEPC, showing relative ASCL1^high^/FOXA2^low^ and ASCL1^low^/FOXA2^high^ patterns, respectively. These categories indicate relative expression dominance rather than mutually exclusive lineages.

To define cluster 13, we prioritized transcription factors using four criteria: higher expression than in adenocarcinoma clusters, higher expression than in cluster 11, higher inferred activity than in cluster 11, and concordant upregulation in clinical ASCL1^low^ versus ASCL1^high^ NEPC samples. FOXA2 was the only TF that met all four criteria, supporting cluster 13 as a FOXA2^high^/ASCL1^low^ terminal NEPC state (Fig. 3B,C; Supplementary Table S3). We then asked whether similar ASCL1/FOXA2 heterogeneity was present in human disease. In the published clinical scRNA-seq dataset, ASCL1^low^ NEPC cells showed higher FOXA2 expression than ASCL1^high^ cells (Fig. 3D) [29]. IHC further identified spatially distinct ASCL1^high^/FOXA2^low^ and ASCL1^low^/FOXA2^high^ regions within the same tumor in 9 of 10 clinical NEPC samples (Fig. 3E).

The two terminal NEPC states also showed different temporal distributions. Cluster 11 emerged earlier during post-castration progression, whereas cluster 13 became prominent later and was enriched in fully relapsed tumors (Fig. 2A,B). Consistent with this pattern, ASCL1^+^/FOXA2^-^cells predominated during early NEPC emergence at 12 and 16 weeks post castration, whereas FOXA2-expressing populations, including ASCL1^+^/FOXA2^+^ and ASCL1^-^/FOXA2^+^, became more prominent in fully relapsed tumors (Fig. 3F). IHC comparing early NE transition with fully relapsed tumors showed a comparable change in ASCL1/FOXA2 expression patterns (Fig. 3G).

Gene set enrichment analysis revealed additional biological differences between clusters 11 and 13. The FOXA2^high^ state was enriched for ribosome biogenesis pathways relative to the ASCL1^high^ state, suggesting stronger ribosome-biogenesis activity (Supplementary Table S4). Together, these findings identify at least two transcriptionally distinct terminal NEPC states with different temporal distributions in the LTL331/331R model, with corresponding ASCL1/FOXA2 expression patterns observed in clinical NEPC.

### 4. RUNX1T1 is induced early and sustained across terminal NEPC states

Having defined an intermediate state and two transcriptionally distinct terminal NEPC states, we next sought regulators induced in the intermediate state and retained across both terminal NEPC states. We examined 2,215 transcriptional regulators represented in the LTL331/331R single-cell dataset [30, 31].

Comparison of terminal NEPC clusters with adenocarcinoma clusters identified 55 significantly altered transcriptional regulators (|fold change|>2, P<0.001), including established NEPC-associated factors such as SOX2, FOXA2, NKX2-1, INSM1, and ASCL1 (Fig. 4A, B). Comparison of cluster 16 with adenocarcinoma clusters identified 30 altered transcriptional regulators (Fig. 4A,C). Fourteen changed concordantly in cluster 16 and terminal NEPC, including six upregulated and eight downregulated genes (Fig. 4A).

**Figure 4.**
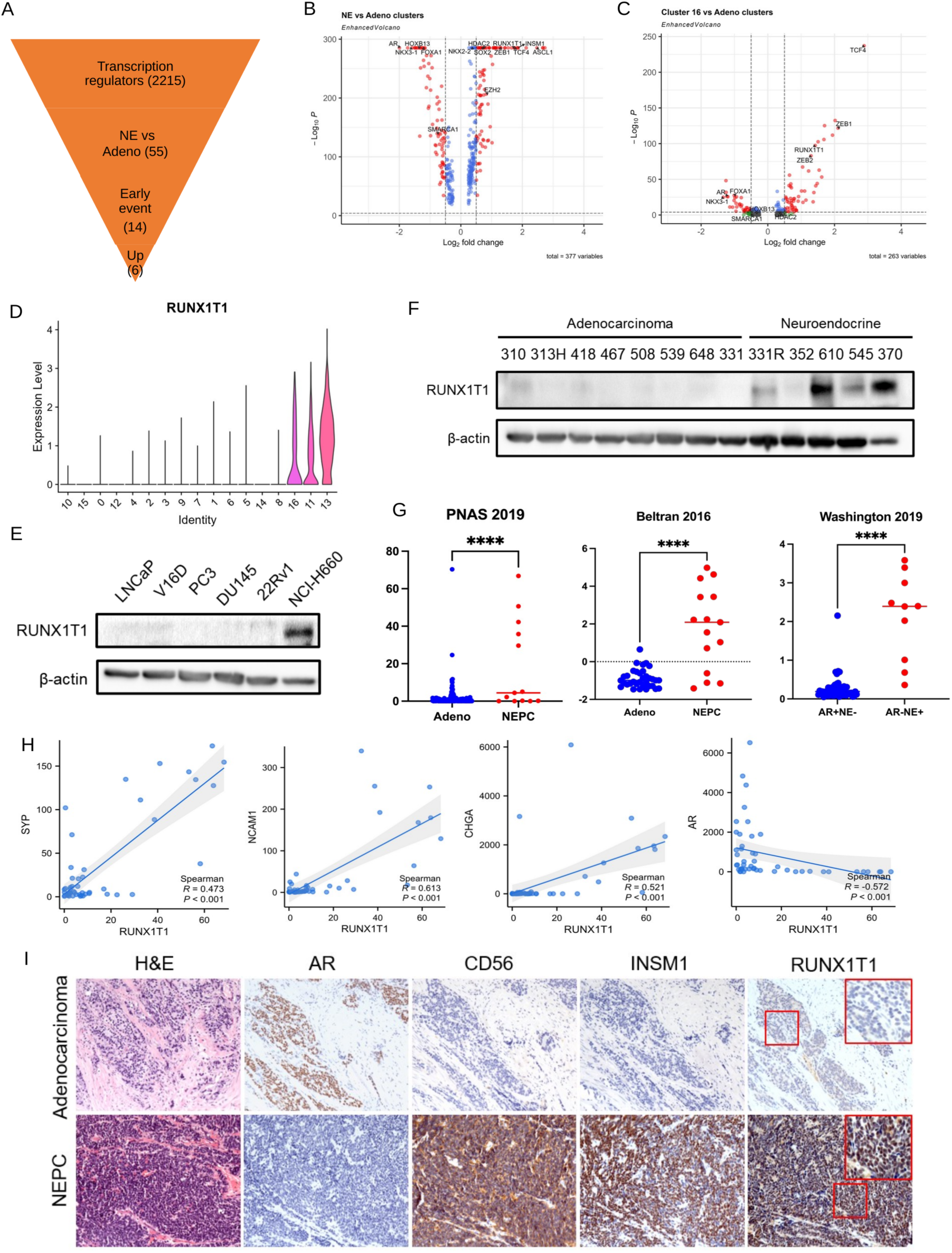
RUNX1T1 is induced early and sustained across terminal NEPC states. **(A)** Stepwise prioritization of transcriptional regulators. Among 2,215 regulators screened, 55 met the differential-expression criteria in terminal NEPC clusters 11 and 13 relative to adenocarcinoma clusters, 14 changed concordantly in cluster 16 and terminal NEPC, and six were concordantly upregulated. **(B,C)** Volcano plots comparing terminal NEPC clusters 11 and 13 with adenocarcinoma clusters (B) and cluster 16 with adenocarcinoma clusters (C); 30 regulators met the differential-expression criteria in the latter comparison. The x-axis shows log₂ fold change, the y-axis shows -log₁₀ P, point colors indicate threshold categories, dashed lines indicate the fold-change and P-value cutoffs, and selected lineage-associated regulators are labeled. **(D)** Violin plot of RUNX1T1 expression across tumor-cell clusters. **(E)** RUNX1T1 immunoblotting in adenocarcinoma cell lines (LNCaP, V16D, PC3, DU145, 22Rv1) and NEPC cell line NCI-H660. **(F)** RUNX1T1 immunoblotting across adenocarcinoma and NEPC PDX models. β-actin was used as the loading control in E and F. **(G)** RUNX1T1 expression in adenocarcinoma and NEPC samples from the Abida et al. (2019) and Beltran et al. (2016) cohorts and in AR^+^/NE^-^and AR^-^/NE^+^ samples from the Labrecque et al. (2019) cohort [32–34]. Horizontal lines indicate medians; four asterisks denote P < 0.0001. **(H)** Correlations between RUNX1T1 and SYP, NCAM1, CHGA, or AR expression in clinical prostate cancer samples. Spearman correlation coefficients and P values are shown. **(I)** Representative hematoxylin and eosin (H&E) staining and IHC for AR, CD56, INSM1, and RUNX1T1 in clinical adenocarcinoma and NEPC specimens.

The concordantly downregulated genes included the luminal lineage regulators AR, FOXA1, GATA2, and NKX3-1, consistent with loss of adenocarcinoma identity during lineage transition. By contrast, several established NEPC regulators, including SOX2, ASCL1, FOXA2, NKX2-1, and INSM1, were altered primarily in terminal NEPC and not in cluster 16 (Fig. 4B,C). We therefore prioritized regulators already induced in cluster 16 and retained in terminal NEPC, a temporal pattern consistent with potential roles in both the acquisition and maintenance of NE-associated features. Among the upregulated candidates, RUNX1T1 was strongly induced in cluster 16 and remained elevated in both terminal NEPC clusters, leading us to prioritize it for functional investigation (Fig. 4D).

RUNX1T1 expression was higher in the NEPC cell line NCI-H660 and in multiple NEPC PDX models than in adenocarcinoma models (Fig. 4E,F). Across three clinical cohorts, RUNX1T1 was consistently increased in NEPC and positively correlated with NE lineage markers (Fig. 4G,H) [32–34]. IHC further confirmed increased RUNX1T1 protein in clinical NEPC specimens (Fig. 4I). Together, these findings identify RUNX1T1 as an early-induced and sustained candidate regulator of NEPC lineage transition and maintenance.

### 5. RUNX1T1 enhances ARPI-induced neuroendocrine features and cell viability in prostate adenocarcinoma cells

Because RUNX1T1 was induced in the early intermediate state, we tested whether it could enhance NE-associated changes in adenocarcinoma cells during AR pathway inhibition. RUNX1T1 was stably overexpressed in V16D cells, an AR^+^/NE^-^/RUNX1T1^-^castration-resistant prostate cancer (CRPC) cell line, and its effects were assessed under baseline conditions and during enzalutamide treatment (Fig. 5A). Under serum-containing conditions, RUNX1T1 overexpression produced only a modest increase in NCAM1 (Fig. 5B,C). Following enzalutamide treatment, however, RUNX1T1-overexpressing cells showed a marked increase in NCAM1 compared with control cells (Fig. 5B,C). qPCR and immunoblotting confirmed increased NCAM1 expression and showed induction of additional NE markers together with reduced AR-lineage output (Fig. 5D,E). These findings indicate that RUNX1T1 overexpression enhances NE-associated changes during AR pathway inhibition.

**Figure 5.**
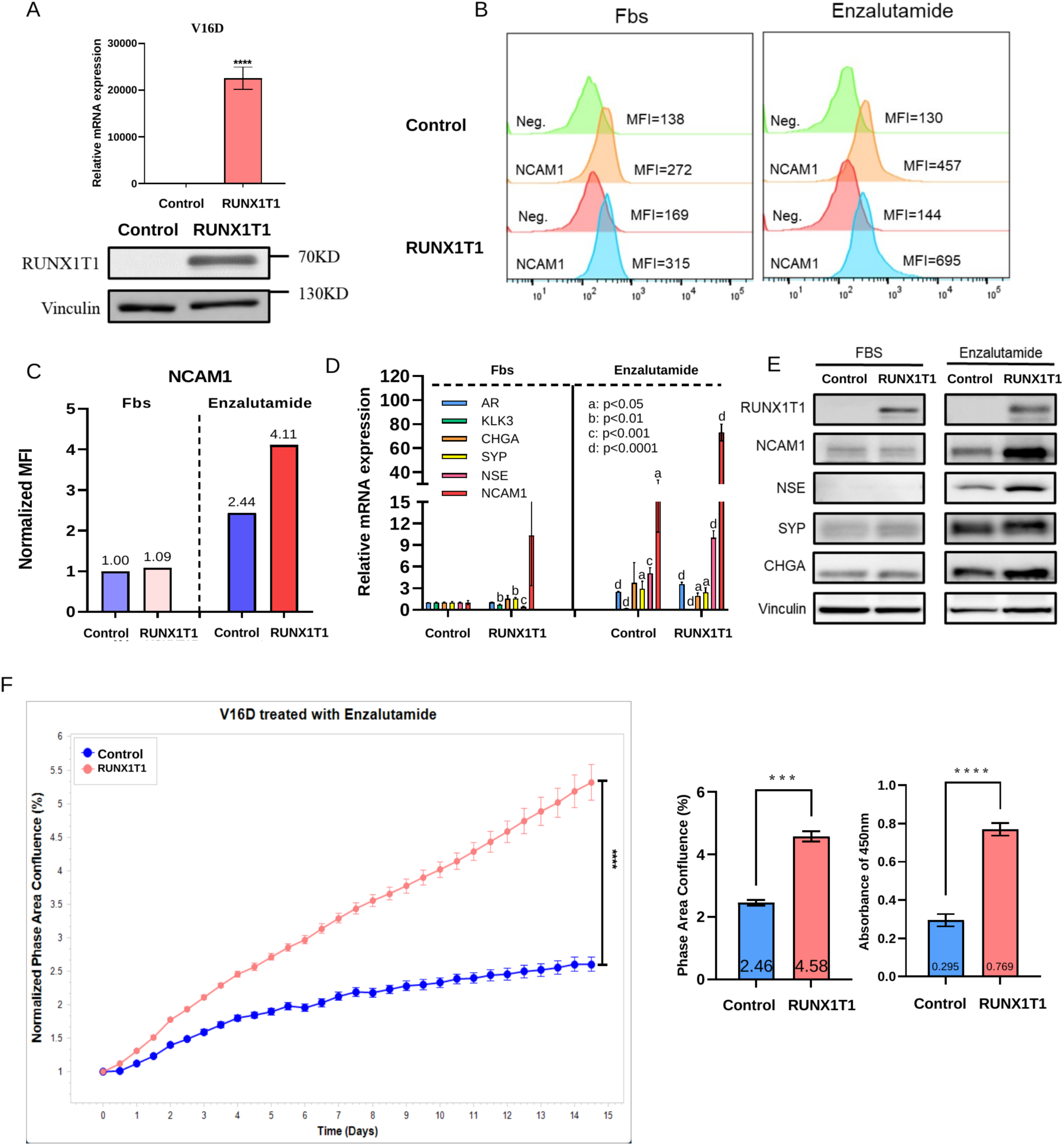
RUNX1T1 enhances NE-associated changes and cell viability during AR pathway inhibition. **(A)** RT-qPCR and immunoblot confirmation of stable RUNX1T1 overexpression in vector-control and RUNX1T1-overexpressing V16D cells. Vinculin was used as the loading control. **(B)** Flow-cytometry histograms of surface NCAM1 in control and RUNX1T1-overexpressing V16D cells cultured in FBS-containing medium or under AR pathway inhibition with 10% charcoal-stripped serum and 10 μM enzalutamide for 2 weeks. Negative-control profiles and mean fluorescence intensity (MFI) values are shown. **(C)** NCAM1 MFI normalized to the vector-control FBS condition. **(D)** Relative mRNA expression of the AR-lineage genes AR and KLK3 and the NE-associated genes CHGA, SYP, NSE, and NCAM1 under the indicated conditions. The letters a, b, c, and d denote P < 0.05, P < 0.01, P < 0.001, and P < 0.0001, respectively. **(E)** Immunoblotting of RUNX1T1, NCAM1, NSE, SYP, and CHGA under the indicated conditions; vinculin was used as the loading control. **(F)** Normalized live-cell phase-area confluence over 15 days during treatment with 10 μM enzalutamide, with control and RUNX1T1-overexpressing cells shown in blue and red, respectively. Day-15 phase-area confluence and CCK-8 absorbance at 450 nm are shown as separate endpoint readouts. Data are mean ± SEM; three and four asterisks denote P < 0.001 and P < 0.0001, respectively.

Given the association between NE lineage plasticity and resistance to AR pathway inhibition, we next tested whether RUNX1T1 affected cell viability during enzalutamide treatment. RUNX1T1-overexpressing cells showed significantly greater cell viability during enzalutamide treatment than control cells (Fig. 5F). Together, these results indicate that RUNX1T1 enhances ARPI-induced NE-associated features and increases cell viability during AR pathway inhibition in adenocarcinoma cells.

### 6. RUNX1T1 supports maintenance of terminal NEPC identity, proliferation, and survival

Because RUNX1T1 remained elevated across both terminal NEPC states, we next tested whether it contributes to maintenance of the established NEPC phenotype. We generated inducible RUNX1T1 knockdown in NCI-H660 NEPC cells using two independent shRNAs. RUNX1T1 knockdown reduced the NE markers NCAM1 and NSE as well as ASCL1 and FOXA2, the transcription factors preferentially associated with clusters 11 and 13 (Fig. 6A,B). These changes indicate disruption of NE-associated transcriptional programs.

**Figure 6.**
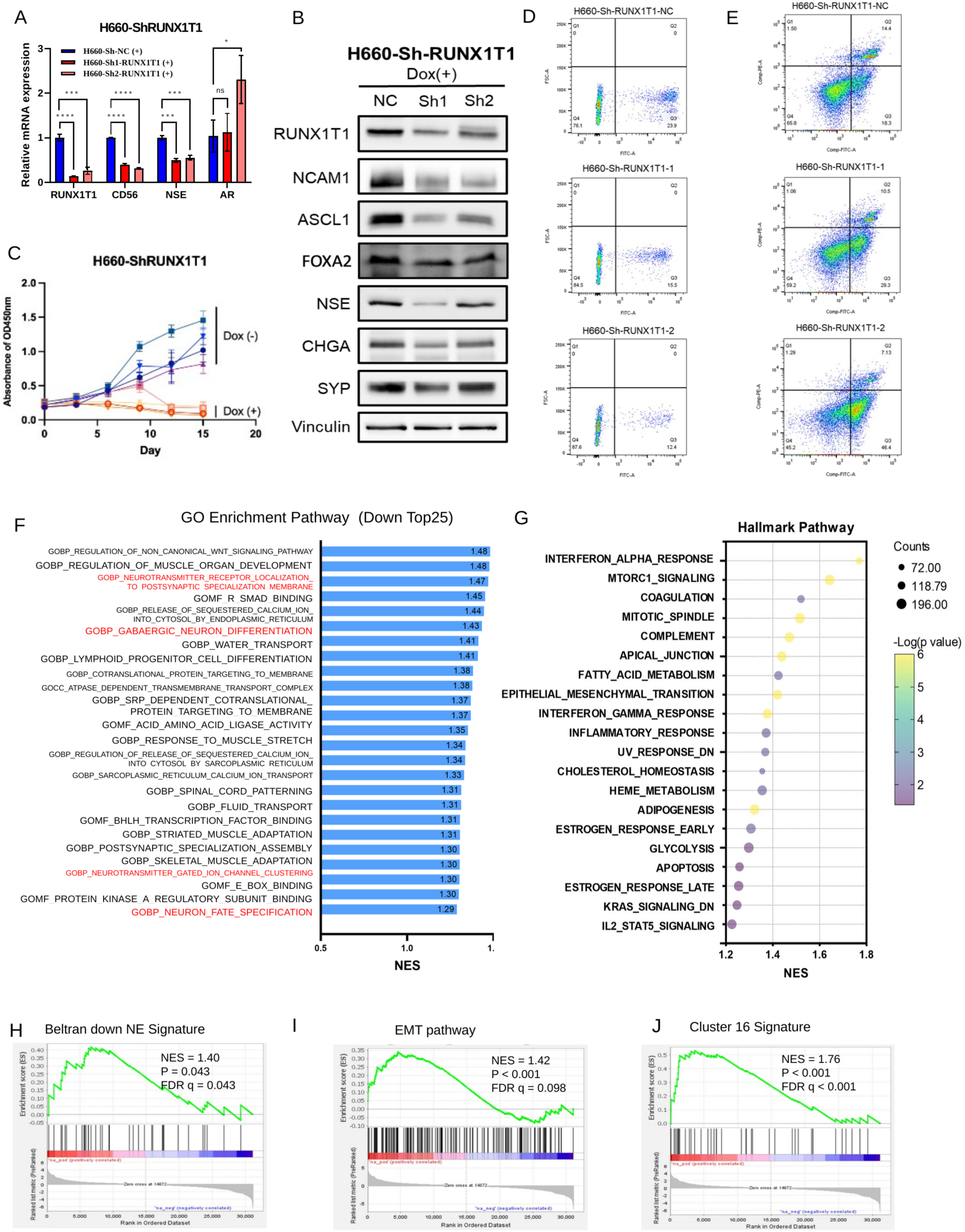
RUNX1T1 knockdown disrupts NE-associated programs and reduces proliferation and survival in NCI-H660 cells. **(A)** RT-qPCR analysis of RUNX1T1, NCAM1/CD56, NSE, and AR in NCI-H660 cells expressing non-targeting control shRNA (NC) or either of two doxycycline-inducible RUNX1T1 shRNAs (Sh1 and Sh2) after doxycycline treatment. *P < 0.05, **P < 0.01, ***P < 0.001, ****P < 0.0001 **(B)** Immunoblotting of RUNX1T1, NCAM1, ASCL1, FOXA2, NSE, CHGA, and SYP after doxycycline induction; vinculin was used as the loading control. **(C)** CCK-8 absorbance at 450 nm over 15 days in NC, Sh1, and Sh2 cells cultured with or without doxycycline. Data are mean ± SEM. **(D)** Representative EdU flow-cytometry plots after doxycycline induction. **(E)** Representative Annexin V/PI flow-cytometry plots after doxycycline induction. Percentages within the indicated gates are shown in D and E. **(F)** Top 25 Gene Ontology biological-process terms reduced after RUNX1T1 knockdown; neuronal-development and synaptic terms are highlighted. **(G)** Hallmark programs enriched after RUNX1T1 knockdown. Dot size indicates gene count, and color indicates - log₁₀ P. **(H-J)** Gene set enrichment analysis of the published Beltran NEPC-down signature (H), Hallmark EMT pathway (I), and cluster 16-derived signature (J) in RUNX1T1-knockdown cells. Black vertical bars indicate signature genes, and the green curve indicates the running enrichment score. Positive enrichment indicates enrichment in RUNX1T1-knockdown cells. NES, nominal P values, and FDR q values are shown: H, NES = 1.40, P = 0.043, FDR q = 0.043; I, NES = 1.42, P < 0.001, FDR q = 0.098; and J, NES = 1.76, P < 0.001, FDR q < 0.001.

RUNX1T1 knockdown also reduced cell growth over time (Fig. 6C). To determine the basis for this effect, we assessed proliferation and apoptosis. EdU incorporation decreased after RUNX1T1 knockdown, indicating fewer cells in S phase (Fig. 6D). Annexin V/PI staining showed increased apoptosis (Fig. 6E). Together, these findings indicate that RUNX1T1 supports NCI-H660 cell growth through effects on both proliferation and survival.

To define the broader transcriptional consequences of RUNX1T1 knockdown, we performed RNA-seq and pathway analysis in control and RUNX1T1-knockdown NCI-H660 cells. Neuronal development and NEPC-associated transcriptional programs were broadly reduced after RUNX1T1 knockdown (Fig. 6F,H; Supplementary Table S5). Conversely, Hallmark pathway analysis showed enrichment of EMT-associated programs following RUNX1T1 knockdown (Fig. 6G,I, Supplementary Tables S5 and S6). More broadly, a cluster 16-derived gene signature was enriched in RUNX1T1-knockdown cells, suggesting a partial transcriptional shift away from terminal NEPC toward the putative intermediate state represented by cluster 16 (Fig. 6J, Supplementary Tables S5 and S6). Together, these findings indicate that RUNX1T1 supports maintenance of the established terminal NEPC phenotype, proliferation, and survival in NCI-H660 cells.

### 7. RUNX1T1 associates with chromatin-associated repressor complexes that include G9A and LASP1 in NEPC cells

RUNX1T1 functions as a transcriptional corepressor through multiprotein complexes. To map its chromatin-associated interactome in NEPC, we performed modified RIME using FLAG immunoprecipitation in NCI-H660 cells stably expressing FLAG-tagged RUNX1T1 (Fig. 7A). Proteins were retained only if enriched relative to both OE-IgG and Control-FLAG backgrounds, reducing nonspecific antibody-and FLAG-associated signals. This dual-background analysis identified 1,058 candidate RUNX1T1-associated proteins (Fig. 7C).

**Figure 7.**
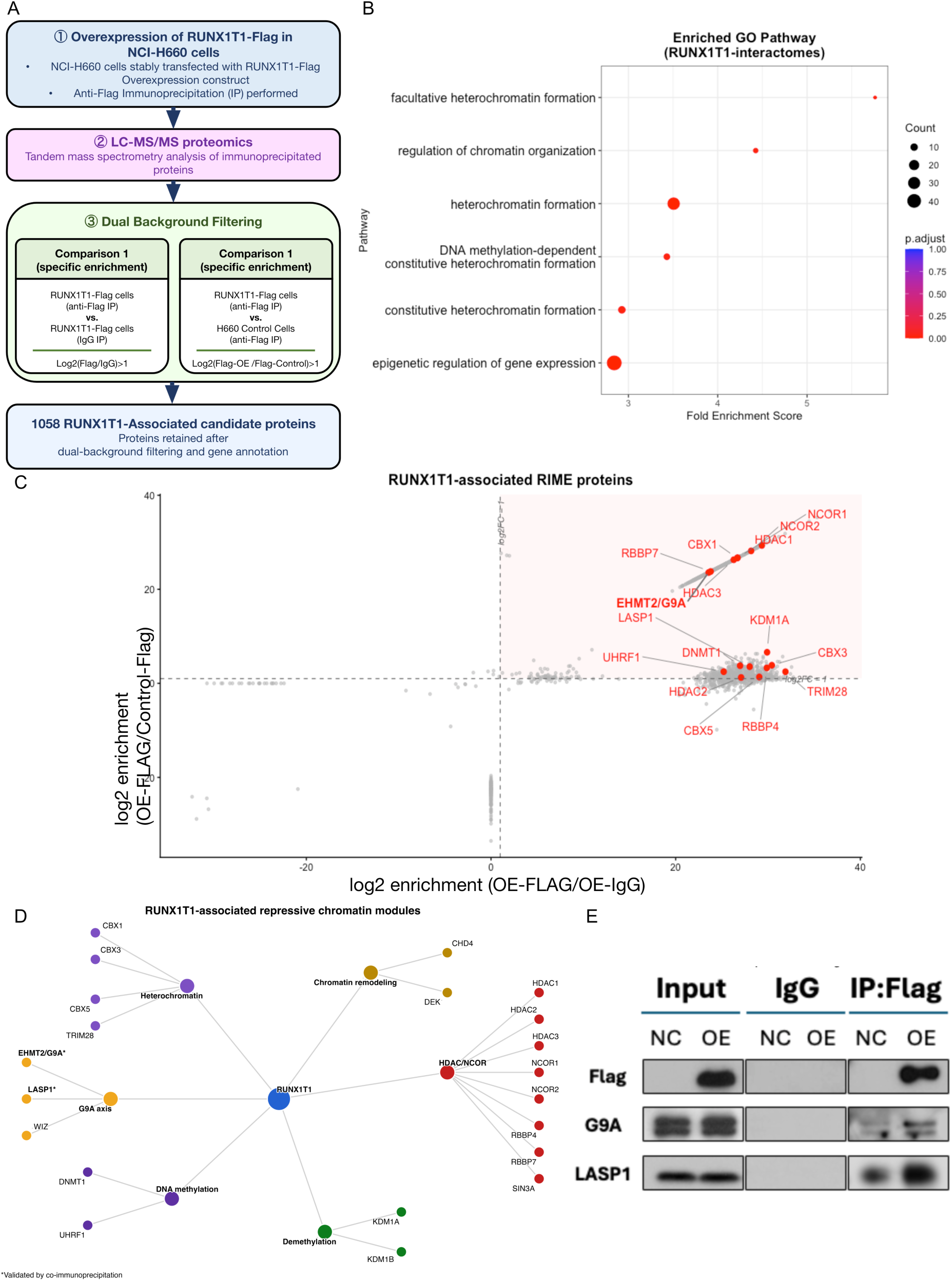
RUNX1T1 associates with chromatin-repressive machinery that includes G9A and LASP1 in NCI-H660 cells. **(A)** Modified rapid immunoprecipitation mass spectrometry of endogenous proteins (RIME) workflow in NCI-H660 cells stably expressing FLAG-tagged RUNX1T1. Anti-FLAG pulldowns from RUNX1T1-FLAG cells were compared with IgG pulldowns from the same cells and anti-FLAG pulldowns from control cells. Proteins with log₂ enrichment greater than 1 in both comparisons were retained, yielding 1,058 candidate RUNX1T1-associated proteins. **(B)** Gene Ontology biological-process enrichment among candidate proteins. Dot size indicates protein count, color indicates adjusted P value, and the x-axis indicates fold-enrichment score. **(C)** Dual-background enrichment plot. The x-axis shows log₂ enrichment in RUNX1T1-FLAG anti-FLAG relative to RUNX1T1-FLAG IgG pulldowns, and the y-axis shows log₂ enrichment in RUNX1T1-FLAG anti-FLAG relative to control-cell anti-FLAG pulldowns. Dashed lines indicate the log₂-enrichment threshold of 1; proteins passing both filters are red, and selected chromatin-associated candidates are labeled. **(D)** Candidate proteins organized by annotated chromatin-regulatory functions. Node colors indicate functional modules, and edges indicate functional grouping and RIME association rather than direct protein-protein binding. Asterisks identify G9A/EHMT2 and LASP1, which were examined by co-immunoprecipitation. **(E)** FLAG co-immunoprecipitation using control (NC) and RUNX1T1-FLAG-overexpressing (OE) NCI-H660 cells. Input, IgG, and anti-FLAG precipitates are shown; G9A and LASP1 were detected in anti-FLAG precipitates from RUNX1T1-FLAG cells.

Functional annotation of these candidates revealed enrichment for chromatin regulation and epigenetic repression, including heterochromatin formation, chromatin organization, DNA methylation-dependent heterochromatin formation, and epigenetic regulation of gene expression (Fig. 7B; Supplementary Table S7). Consistent with this enrichment, RUNX1T1-associated proteins included EHMT2/G9A, HP1 family proteins, UHRF1, KDM1A/LSD1, and HDAC/corepressor components (Fig. 7C,D; Supplementary Table S7). Among these candidates, EHMT2/G9A was strongly enriched, and LASP1 was also recovered. Because a LASP1-G9A repressive complex was previously characterized during NEPC progression in this model, recovery of both proteins suggested that RUNX1T1 may associate with this epigenetic axis [27].

Co-immunoprecipitation in RUNX1T1-FLAG NCI-H660 cells confirmed that both EHMT2/G9A and LASP1 were present in RUNX1T1-containing complexes (Fig. 7E). Together, these findings indicate that RUNX1T1 associates with G9A/LASP1-containing chromatin repressor complexes, suggesting a mechanistic link between RUNX1T1 and chromatin repression in NEPC.

## Discussion

t-NEPC is a lethal manifestation of lineage plasticity, but the tumor-cell states connecting treated adenocarcinoma with relapsed NEPC remain incompletely defined. Genetically engineered models have shown that defined genetic alterations can produce NEPC-like states [16, 18, 35–38], whereas longitudinal human-derived systems that capture progression under AR pathway suppression remain limited. The LTL331/331R model undergoes adenocarcinoma-to-NEPC progression after host castration and recapitulates the donor patient’s clinical course [20–22]. Analysis across this time course resolved an AR^low^/NE^low^ intermediate state after castration and two terminal NEPC states distinguished by relative ASCL1 and FOXA2 programs. Integrating this temporal framework with functional studies identified RUNX1T1 as a regulator induced before overt relapse and retained after NEPC was established. These findings connect early treatment-associated reprogramming with terminal-state heterogeneity and maintenance of NEPC.

Cluster 16 corresponded to the EMT-enriched differentiation node previously reported in this model [27]. It appeared after castration and before terminal NEPC became dominant, occupied intermediate positions in trajectory analyses, and combined attenuated adenocarcinoma programs with incomplete NE activation. Its enrichment for EMT, stem cell-associated, metabolic, and chromatin-regulatory programs is consistent with a lineage-plastic transitional state. This state provides a biologically informative window in which regulators induced during early remodeling can be distinguished from factors activated only after terminal NEPC is established. However, temporal sampling and trajectory inference do not establish ancestry. In the absence of direct lineage tracing, cluster 16 remains a putative transitional state, and alternative trajectories cannot be excluded. Longitudinal clinical sampling will be required to determine whether comparable states occur in patients and whether they predict progression to NEPC.

Terminal NEPC was also transcriptionally heterogeneous. Clusters 11 and 13 differed in relative ASCL1 and FOXA2 expression and activity, temporal distribution, and broader transcriptional programs. ASCL1^+^/FOXA2^-^cells predominated during early NEPC emergence, whereas FOXA2-expressing populations, including ASCL1^+^/FOXA2^+^ and ASCL1^-^/FOXA2^+^ cells, became more prominent in fully relapsed tumors. Corresponding expression relationships in published clinical scRNA-seq data and spatially distinct patterns within clinical tumors support the relevance of this heterogeneity beyond the PDX model [29]. This temporal shift is compatible with some terminal heterogeneity developing during progression rather than reflecting only stable, parallel identities, consistent with dynamic ASCL1-dependent NEPC evolution in another model [19]. Importantly, these states reflect relative transcriptional dominance rather than mutually exclusive lineages. The presence of double-positive cells in relapsed tumors indicates that their relative activities may distinguish intratumoral states even when both factors are coexpressed. Longitudinal clinical single-cell and spatial studies will be needed to establish the prevalence, stability, interconversion, temporal ordering, and therapeutic vulnerabilities of these states.

Within this temporal and heterogeneous framework, RUNX1T1 was notable because it was induced before overt NEPC and retained across both terminal programs. This early and sustained pattern is consistent with a regulator that may contribute to the acquisition of NE-associated features and remain relevant to their maintenance after NEPC is established. RUNX1T1 is an MTG-family transcriptional corepressor that functions through multiprotein repressor complexes [39–43] and has been implicated in several solid tumors [44–46]. Recent work established RUNX1T1 as a cross-cancer regulator of lineage plasticity and identified an NCOR/HDAC3-dependent chromatin mechanism [47]. Its cross-cancer design provided broad mechanistic and therapeutic insight but did not resolve when RUNX1T1 is induced during treatment-induced adenocarcinoma-to-NEPC progression or whether it remains relevant across heterogeneous terminal NEPC states. Our findings complement this work by placing RUNX1T1 induction in the AR-low/NE-low intermediate state, showing its persistence across ASCL1-and FOXA2-associated terminal programs, and linking it to G9A/LASP1-containing chromatin repressor complexes in NEPC. Its enrichment in independent NEPC models and clinical cohorts further supports its relevance beyond LTL331/331R.

Functional studies were consistent with roles at both stages. In V16D adenocarcinoma cells, RUNX1T1 overexpression enhanced NE-associated changes and increased cell viability during enzalutamide treatment, supporting a contribution to treatment-associated lineage remodeling. RUNX1T1 remained elevated across both terminal NEPC states, and its knockdown reduced both ASCL1 and FOXA2, consistent with relevance across these distinct transcriptional programs. In NCI-H660 cells, RUNX1T1 knockdown also suppressed broader neuronal and NEPC-associated transcriptional programs and reduced proliferation and survival. Enrichment of a cluster 16-derived signature after knockdown further suggested a partial transcriptional shift away from terminal NEPC toward an intermediate-like program. Together, these complementary gain-and loss-of-function findings support a role for RUNX1T1 in both the acquisition of NE-associated features during AR pathway inhibition and their maintenance after NEPC is established.

The chromatin interactome provides mechanistic context for these phenotypes. RIME identified candidate associations with G9A, LASP1, HP1 family proteins, UHRF1, LSD1, and HDAC-related corepressors, and co-immunoprecipitation confirmed that G9A and LASP1 were present in RUNX1T1-containing complexes. Identification of G9A and LASP1 among the RIME candidates links RUNX1T1 to a repressive axis previously implicated in NEPC progression in this model [27], whereas the HDAC-related associations are consistent with recent evidence for RUNX1T1-NCOR/HDAC3-dependent chromatin reprogramming [47]. The identification of LSD1 is also consistent with its reported role in NEPC cell survival through noncatalytic TP53 repression in cooperation with HDAC2 [48]. Rather than defining a linear RUNX1T1-G9A-LASP1 pathway, these results place RUNX1T1 within a broader chromatin-repressive network in NEPC. Mapping RUNX1T1 genomic occupancy and testing whether G9A, LASP1, or HDAC components are required for its transcriptional effects will be necessary to establish the operative mechanism.

Several limitations define the scope of these conclusions. The temporal framework derives from a single PDX model without direct lineage tracing, precluding proof that cluster 16 directly gives rise to the terminal states or exclusion of additional transition routes. Clinical specimens support ASCL1/FOXA2 heterogeneity and RUNX1T1 expression but do not establish temporal ordering in patients. Functional studies centered on one established NEPC cell line, and the direct genomic targets and partner dependencies of RUNX1T1 remain unresolved. Despite these limitations, the integrated longitudinal, functional, and clinical evidence resolves a treatment-associated intermediate state, reveals temporally changing heterogeneity within terminal NEPC, and identifies RUNX1T1 as an early-induced and sustained regulator linked to both lineage transition and maintenance. These findings connect early therapy-associated reprogramming with the biology of established NEPC and provide a framework for investigating regulatory dependencies that persist across disease progression.

## Acknowledgments

This research was supported in part by the Canadian Institutes of Health Research (#180554, #186331, #205704, #205896, and #212791), Terry Fox Research Institute (#1109), a Canadian Cancer Society Breakthrough Team Grant generously supported by the Lotte & John Hecht Memorial Foundation (CCS grant #707683), the BC Cancer Foundation (grant #1PRRG012), and support from the China Scholarships Council (to Y.N.; No. 202206240124).

## Author contributions

**Y.N.**, **M.S.**, **D.L.**, **H.Z.**, **C.C.**, and **YZ.W.** conceptualized and designed the study. **Y.N.**, **M.S., L.L., and W.D.** performed the experiments. **H.X.**, **D.L.**, and **X.D.** contributed to PDX model development and pathological analysis. **R.W.** performed immunohistochemical staining. **A.H.** performed sequencing experiments. **D.L.**, **Y.Y.L.**, **T.M.**, **R.B.**, **X.P.**, **A.C., Y.N.,** and **M.S.** contributed to data analysis and figure generation. **S.L.B.** contributed to project management. **F.S.**, **Yu.W**., **J.C.**, **N.X.**, **N.L.**, **M.G.**, and **C.O.** contributed to scientific discussion and data interpretation. **G.W.** and **V.W.** provided patient tumor specimens. **Y.N.**, **M.S.**, **D.L.**, and **YZ.W.** wrote the manuscript with input from all authors. **H.Z.**, **C.C.**, and **YZ.W.** jointly supervised the study. All authors read and approved the final manuscript.

## Data Availability Statement

The published clinical datasets analyzed in this study are available through the original studies cited [29, 32–34]. The longitudinal LTL331/331R PDX single-cell RNA-sequencing data analyzed in this study are available under NCBI BioProject PRJNA1263525 [27]. The shared shNC bulk RNA-sequencing control data are available in the NCBI Gene Expression Omnibus under accession GSE277727 [14]. The shRUNX1T1 bulk RNA-sequencing data generated in this study are being prepared for deposition in GEO, and the accession number will be added upon assignment.

## Methods

### Ethics Statement

Human prostate cancer specimens used in this study were obtained with written informed consent through the British Columbia Genitourinary Cancer Biobank under protocols approved by the University of British Columbia Clinical Research Ethics Board and affiliated hospital research ethics committees (certificates H21-03722 and H24-00244). All specimens were de-identified before analysis, and the human research was conducted in accordance with institutional guidelines and the Declaration of Helsinki.

The animal procedures used to generate the LTL331/331R tissues analyzed in this study, including host castration and tumor collection, were performed in accordance with the guidelines of the Canadian Council on Animal Care under protocols approved by the University of British Columbia Animal Care Committee (certificates A20-0221, A24-0218, and A24-0165).

### PDX models and clinical datasets

LTL331 PDX models were established as previously described [20]. After host castration, PDX tissue samples were harvested at defined time points spanning pre-castration disease, post-castration regression, and relapsed NEPC. Xenograft tissues were measured, fixed for histopathology, and processed for downstream molecular analyses, including single-cell RNA sequencing. Publicly available and in-house clinical datasets were used to validate key transcriptional states and marker relationships, as described in the Results and figure legends.

### Single-cell RNA sequencing and bioinformatic analysis

The LTL331/331R single-cell RNA-seq dataset was previously generated and reported by our group [27] (BioProject PRJNA1263525). In the present study, raw sequencing data were reprocessed and reanalyzed to examine intermediate tumor-cell states, terminal NEPC heterogeneity, and RUNX1T1-associated regulatory programs. The main analysis included seven time-course samples spanning pre-castration adenocarcinoma, post-castration regression, and relapsed NEPC; the 22-week time point included in the prior report was excluded from the present analysis to avoid disproportionate representation of the fully relapsed state in clustering and trajectory analyses. Cell numbers per time point are provided in Supplementary Table S1. Clustering, lineage scoring, trajectory inference, and pathway analyses were performed as described below.

### Read alignment and human tumor-cell selection

Raw single-cell RNA sequencing reads from the LTL331/331R time-course samples were processed using Cell Ranger. Reads were aligned to a combined human-mouse reference genome generated from Ensembl Release 90 GRCh38 human genome annotation and GRCm38 mouse genome annotation. To distinguish human tumor cells from mouse stromal cells in PDX samples, cells with more than 80% of reads uniquely mapped to the human genome were retained for downstream analysis. Human-specific reads were used to generate filtered count matrices, and samples were aggregated into a single count matrix using cellranger aggr without normalization.

### Quality control and clustering

The aggregated count matrix was analyzed using Seurat v4.1.1. Genes detected in fewer than 10 cells were removed. Cells were retained if they had mitochondrial transcript percentages below 25%, at least 1,000 UMIs, and log-transformed UMI counts and detected gene numbers within three median absolute deviations from the median. Data were log-normalized, variable features were selected, and gene expression values were scaled using default Seurat parameters. Principal component analysis was performed, and the top 50 principal components were selected based on the knee point of the variance distribution. Cells were clustered using the FindNeighbors and FindClusters functions and visualized by UMAP.

### Trajectory inference

Relationships among tumor-cell states were inferred using partition-based graph abstraction (PAGA), implemented in Scanpy v1.10.1, and Palantir v1.3.3. PAGA was applied to the nearest-neighbor graph using the final tumor-cell cluster assignments to estimate connectivity between clusters. Palantir was independently applied to the tumor-cell dataset using pre-castration cluster 10 as the origin to calculate pseudotime across the treatment-associated cell states.

### Single-cell regulatory network analysis

Single-cell regulatory network inference was performed using pySCENIC v0.12.0 to identify active regulons across tumor-cell states. To reduce computational burden, 7,878 cells were subsampled from the full dataset, with up to 500 cells included per cluster. Because pySCENIC is nondeterministic, the analysis was repeated 20 times. Transcription factors detected in at least 16 of 20 runs were retained, and target genes were included in the final regulon only if they were identified in at least 80% of runs in which the corresponding transcription factor was detected. This analysis identified 200 active regulons. Regulon activity was quantified by AUC scores, and unsupervised clustering based on regulons ranked among the top five in at least one cluster was used to compare adenocarcinoma, transitional, and NEPC-associated cell states.

### Cell culture and treatment

293T, LNCaP, PC3, DU145, 22Rv1, and NCI-H660 cells were obtained from the American Type Culture Collection (ATCC). Cells were authenticated by fingerprinting at Fred Hutchinson Cancer Research Center. V16D cells were provided by Dr. Amina Zoubeidi. Mycoplasma testing was performed routinely at the Vancouver Prostate Centre. NCI-H660 cells were cultured in RPMI-1640 medium (Gibco) supplemented with 5% fetal bovine serum (FBS; Gibco), 10 nmol/L β-estradiol (Sigma), 10 nmol/L hydrocortisone (Sigma), and 1% insulin-transferrin-selenium (Thermo Fisher). LNCaP, PC3, DU145, 22Rv1, and V16D cells were maintained in RPMI-1640 containing 10% FBS. 293T cells were maintained in DMEM (Gibco) containing 10% FBS. For in vitro induction of neuroendocrine features in V16D cells, cells were cultured for 24 h in phenol red-free RPMI-1640 containing 10% charcoal-stripped serum and then treated with 10 μM enzalutamide in the same medium for 2 weeks.

### Immunohistochemistry

Formalin-fixed, paraffin-embedded tissue sections were cut at 5 μm, deparaffinized in xylene, and rehydrated through graded ethanol. Antigen retrieval was performed in 10 mM citrate buffer (pH 6.0) using a pressure cooker for 10 minutes. After cooling to room temperature, endogenous peroxidase activity was quenched with 3% hydrogen peroxide for 10 minutes, and sections were blocked in 5% normal goat serum for 30 minutes before overnight incubation with primary antibodies at 4°C. Primary antibodies included anti-RUNX1T1 (Proteintech, #15494-1-AP), anti-NCAM1/CD56 (Cell Marque, #156R), anti-AR (Cell Signaling Technology, #5153S), and anti-INSM1 (Sigma-Aldrich, #MRQ-70).

### Western blotting

Protein lysates were resolved by SDS-PAGE, transferred to PVDF membranes, and blocked in TBST containing 5% skim milk for 1 hour at room temperature. Membranes were incubated with primary antibodies against RUNX1T1 (Proteintech, #15494-1-AP, 1:1000), NCAM1/CD56 (Cell Signaling Technology, #3576, 1:1000), CHGA (Cell Signaling Technology, #60893, 1:1000), SYP (Santa Cruz Biotechnology, #sc-17750, 1:500), ASCL1 (Santa Cruz Biotechnology, #sc-374104, 1:500), FOXA2 (Cell Signaling Technology, #8186, 1:1000), NSE (Cell Signaling Technology, #9536, 1:1000), ACTB/β-actin (Sigma-Aldrich, #A5441, 1:2000), SOX2 (Cell Signaling Technology, #3579, 1:1000), and Vinculin (Sigma-Aldrich, #V9131, 1:2000).

### Plasmids and stable cell line establishment

For overexpression studies, the RUNX1T1 expression plasmid (TFORF1679, Addgene #143835) and corresponding GFP vector control (TFORF3549, Addgene #145025) were used. For knockdown studies in NCI-H660 cells, two shRNA sequences targeting RUNX1T1 were designed: shRNA#1 (CACCGAATGACATTTCACCCGAGAT) and shRNA#2 (CACCGCTGTGCAATACCTTCAAAAA), and cloned into the doxycycline-inducible tet-pLKO-puro plasmid (Addgene #21915).

Lentiviral particles were produced in HEK293T cells by co-transfection of the expression plasmid with the packaging plasmid psPAX2 (Addgene #12260) and the envelope plasmid pMD2.G (Addgene #12259) using Lipofectamine 3000 (Thermo Fisher Scientific, #L3000008) according to the manufacturer’s instructions.

Stable RUNX1T1-overexpressing V16D cells and shRNA-expressing NCI-H660 cells were established by puromycin selection (2 μg/mL for V16D; 1 μg/mL for NCI-H660), with selection maintained during downstream experiments. For knockdown experiments, stable shRNA-expressing NCI-H660 cells were treated with 1 μg/mL doxycycline to induce shRNA expression for the indicated durations.

### RNA Preparation, RT-qPCR, and bulk RNA sequencing

Total RNA was extracted from cells using the RNeasy Plus Mini Kit (Qiagen, #74034) according to the manufacturer’s instructions. One microgram of RNA was reverse transcribed into cDNA using the QuantiTect Reverse Transcription Kit (Qiagen, #205313). Quantitative real-time PCR was performed on an Applied Biosystems QuantStudio 7 Pro instrument and analyzed using Design and Analysis Software version 2.8.0 (Life Technologies). Primer sequences are provided in Supplementary Table S8.

For bulk RNA sequencing, total RNA was extracted from NCI-H660 cells stably expressing shNC or shRUNX1T1 after 8 days of doxycycline treatment. The shNC RNA-seq samples used in the present analysis were previously reported as controls in our study of PROX1 knockdown [14]. The shNC, shPROX1, and shRUNX1T1 samples were submitted for sequencing together. The previous study compared shPROX1 with shNC, whereas the present study compared shRUNX1T1 with the same shNC controls. The shRUNX1T1 samples were not included in the previous publication.

Libraries were prepared and sequenced by Novogene on the Illumina NovaSeq 6000 platform, generating 150-bp paired-end reads at a depth of approximately 20 million reads per sample. Reads from the shNC and shRUNX1T1 samples were processed together and aligned to the GRCh38 reference genome using HISAT2 version 2.0.5. Gene expression was quantified using featureCounts version 1.5.0-p3, and differential expression analysis was performed using DESeq2 version 1.20.0 in R. Genes with an adjusted P value of 0.05 or less were considered differentially expressed. Pathway analysis was performed using Gene Set Enrichment Analysis version 4.3.2 with the EMT and cluster 16-derived gene sets defined in Supplementary Table S6; complete enrichment results are reported in Supplementary Table S5. The shRUNX1T1 RNA-sequencing data generated in this study are being prepared for deposition in the NCBI Gene Expression Omnibus (GEO), and the accession number will be added upon assignment. The shared shNC RNA-sequencing data are available under accession GSE277727.

### Cell viability assay

Cell viability was assessed using the Cell Counting Kit-8 (CCK-8; Dojindo Laboratories, #CK04) according to the manufacturer’s instructions. Cells were seeded in 96-well plates with doxycycline and incubated overnight. At the indicated time points, 20 μL of CCK-8 reagent was added to each well containing 200 μL of medium, and plates were incubated for 3 hours at 37°C in 5% CO₂. Absorbance at 450 nm was then measured.

### Incucyte live-cell imaging

Real-time cell proliferation and confluence were monitored using the Incucyte Live-Cell Analysis System (Sartorius). Cells were seeded in 96-well plates at an optimized density, allowed to adhere overnight, and imaged every 12 hours using a 10× objective. Phase-contrast images were collected from multiple fields per well, and confluence was quantified using the integrated Incucyte analysis software.

### Apoptosis assay

Apoptosis was evaluated using the BD Pharmingen FITC Annexin V Apoptosis Detection Kit I (#556547) according to the manufacturer’s protocol. Cells were harvested after 7 days of doxycycline-induced knockdown and stained with 5 μL of Annexin V and 5 μL of propidium iodide for 15 minutes at room temperature. Samples were analyzed using an LSRFortessa II flow cytometer (BD Biosciences). Unstained controls and single-stained controls were used to establish gating.

### EdU assay

Cell proliferation was assessed by EdU incorporation using the EdU Staining Proliferation Kit (iFluor 488; Abcam #ab219801). Cells were pulsed with 10 μM EdU for 3 hours at the indicated time points, fixed, permeabilized, and stained according to the manufacturer’s instructions. Flow cytometric analysis was performed on an LSRFortessa II instrument (BD Biosciences). Cells not exposed to EdU served as negative controls.

### RIME (Rapid immunoprecipitation mass spectrometry of endogenous proteins)

Chromatin-associated RUNX1T1-interacting proteins were identified using a modified Rapid Immunoprecipitation Mass Spectrometry of Endogenous proteins (RIME) approach [49]. FLAG immunoprecipitation was performed on chromatin from NCI-H660 cells stably expressing FLAG-tagged RUNX1T1, with H660-Control cells serving as a background control and IgG immunoprecipitation as a nonspecific antibody control. Briefly, cells were dual-crosslinked using 2 mM disuccinimidyl glutarate (DSG) for 30 minutes at room temperature, followed by 1% formaldehyde fixation for 10 minutes. Crosslinking was quenched with 125 mM glycine for 5 minutes, and cells were washed twice with ice-cold PBS containing protease inhibitors. Cell pellets were lysed, and chromatin was fragmented by sonication to generate soluble chromatin complexes. Equal amounts of solubilized chromatin were incubated overnight at 4°C with anti-FLAG M2 magnetic beads (Sigma-Aldrich, #M8823) or mouse IgG (Santa Cruz Biotechnology, #sc-2025). Beads were washed extensively under stringent conditions, and bound proteins were digested on-bead with trypsin. Peptides were analyzed by LC-MS/MS. Proteins enriched in H660-RUNX1T1-FLAG anti-FLAG pulldowns relative to both H660-RUNX1T1-FLAG IgG pulldowns and H660-Control anti-FLAG pulldowns were considered candidate RUNX1T1 chromatin interactors.

### Co-immunoprecipitation (co-IP)

For co-immunoprecipitation assays, NCI-H660 cells expressing FLAG-tagged RUNX1T1 were lysed in ice-cold NP-40 lysis buffer supplemented with protease and phosphatase inhibitors. Lysates were clarified by centrifugation, and equal amounts of protein were incubated overnight at 4°C with anti-FLAG magnetic beads. Immunocomplexes were washed extensively, eluted in SDS sample buffer, resolved by SDS-PAGE, and analyzed by immunoblotting using antibodies against RUNX1T1, G9A/EHMT2, LASP1, and other indicated proteins. Input lysates were included as controls. Antibodies used for co-IP included anti-FLAG (Cell Signaling Technology, #14793), anti-G9A (Abcam, #ab185050), and anti-LASP1 (Abclonal, #A3941).

### Statistical analysis

Statistical analyses were performed using GraphPad Prism 10. Comparisons between two groups were assessed using two-tailed Student’s t-tests, and comparisons among three or more groups were assessed by one-way ANOVA followed by Tukey’s post hoc test. Growth curves were analyzed by two-way ANOVA with repeated measures. Spearman correlation coefficients with 95% confidence intervals were used to assess linear relationships between continuous variables. For single-cell RNA-seq differential expression analyses, the Wilcoxon rank-sum test with Benjamini-Hochberg correction was applied, with adjusted P < 0.05 considered significant. For all other analyses, P < 0.05 was considered statistically significant. Significance is indicated as *P < 0.05, **P < 0.01, ***P < 0.001, and ****P < 0.0001. Data are presented as mean ± SEM from at least three independent biological replicates unless otherwise indicated in the figure legends.

